# Hidden population modes in social brain morphology: Its parts are more than its sum

**DOI:** 10.1101/2020.08.07.241497

**Authors:** Hannah Kiesow, R. Nathan Spreng, Avram J. Holmes, M. Mallar Chakravarty, Andre F. Marquand, B.T. Thomas Yeo, Danilo Bzdok

## Abstract

The complexity of social interactions is a defining property of the human species. Many social neuroscience experiments have sought to map ‘perspective taking’, ‘empathy’, and other canonical psychological constructs to distinguishable brain circuits. This predominant research paradigm was seldom complemented by bottom-up studies of the unknown sources of variation that add up to measures of social brain structure; perhaps due to a lack of large population datasets. We aimed at a systematic de-construction of social brain morphology into its elementary building blocks in the UK Biobank cohort (n=~10,000). Coherent patterns of structural co-variation were explored within a recent atlas of social brain locations, enabled through translating autoencoder algorithms from deep learning. The artificial neural networks learned rich subnetwork representations that became apparent from social brain variation at population scale. The learned subnetworks carried essential information about the co-dependence configurations between social brain regions, with the nucleus accumbens, medial prefrontal cortex, and temporoparietal junction embedded at the core. Some of the uncovered subnetworks contributed to predicting examined social traits in general, while other subnetworks helped predict specific facets of social functioning, such as feelings of loneliness. Our population-level evidence indicates that hidden subsystems of the social brain underpin interindividual variation in dissociable aspects of social lifestyle.

## Introduction

Social interaction is a central activity to the human species, which has enabled the construction of civilizations by collaboration across and between generations (Tennie et al., 2009). This realization has led many investigators to adopt the social brain hypothesis (Byrne & Whiten, 1988; Humphrey, 1978). The perspective posits that dimensions of social complexity, like group size (Dunbar & Shultz, 2017; Lewis et al., 2011) or the capacity to anticipate other individuals’ ongoing thought (Powell et al., 2010), have shaped the evolution of brain structure. To this end, the need to adapt to the increasing demands of social complexity and social challenges has likely played a significant role in natural selection, thus influencing the course of primate brain evolution (Byrne & Whiten, 1988; Humphrey, 1978). The significance of social interaction for the human species also becomes apparent in its close relation to mental health. For instance, a lack of regular social interactions is known to escalate the risk for various major psychiatric disorders (Danilo Bzdok & Dunbar, 2020; Cacioppo & Hawkley, 2009; Tost & Meyer-Lindenberg, 2012).

To interrogate the relationship between dimensions of everyday social experience may manifest themselves in the human brain, previous structural brain-imaging studies have established the close relationship between markers of social interaction frequency and quantity and grey matter structure in regions, such as the amygdala (R. Kanai et al., 2012), nucleus accumbens (Kiesow et al., 2020 in press) and ventromedial prefrontal cortex (Dunbar & Shultz, 2017; Lewis et al., 2011). In addition, neuroscientists also commonly rely on carefully curated experimental tasks, which frequently endorse a select set of psychological constructs like ‘theory of mind’, ‘empathy’, or the ‘mirror neuron system’. These hypothesis-guided social, cognitive and affective neuroscience experiments have proven invaluable for localizing neural activity responses in controlled task environments. For instance, in moral decision making, experimental paradigms involving the trolley dilemma have been used to compare the neural correlates underlying emotional-affective processes against those involved in more rational abstract-perspective taking (Danilo Bzdok, Groß, et al., 2015; Sevinc & Spreng, 2014). Recent trends towards large-scale aggregation of social neuroscience experiments have opened the door to principled across-study integration by an arsenal of meta-analysis techniques. These new tools have allowed neuroscientists to identify the parts of the human brain that respond most consistently when participants are engaged in a diverse set of social-affective experiences (Spreng et al. 2009; Alcalá-López et al. 2018; Schurz et al. 2014).

Despite the merits of defining convergence zones related to the social brain based on aggregate summaries from meta-analyses, the constituent regions may obscure distinct social-affective functional systems when collapsing separate studies into averages. In social neuroscience, a majority of previous brain-imaging studies implicitly assumed that a target region is sufficiently described by a single pattern of neural activity obtained through some subtraction analysis, which results in relative increase or decrease of neural response. This pervasive assumption may obfuscate distinct sources of biological variation - *within* trusted convergence zones - that factor into specific effects in a brain region across individuals. As a consequence, much prior social neuroscientific work has been less sensitive to such mutually overlapping sources of variation in the wider population. Yet, adaptive social functioning relies on the dynamic coordination of a host of abilities, ranging from lower-sensory processing of social cues like faces to higher processing such as mental scene construction (Mesulam, 1998).

The current study adopted a data-driven stance on population neuroscience to dissect and characterize separable neural systems from social brain regions that preferentially support social-affective processes. All of our analyses capitalized on the UK Biobank (UKB) - currently the largest, uniformly acquired human brain-imaging dataset in the world - to identify the hidden structural components within the social brain. We further characterized the derived social subnetworks by profiling their predictive role in several social lifestyle markers. Importantly, the overall analytical strategy departs from many previous approaches that assumed each specific brain region underpins a unique element of social functioning. This common a-priori assumption neglects the possible existence of subnetworks that may partially overlap with each other in topography and functional implications across the social brain elements (Shine et al., 2019; R. N. Spreng & Andrews-Hanna, 2015; Yeo et al., 2015). Moreover, many studies restricted themselves to charting patterns in the social brain by clustering or mixture modeling approaches, which strictly assign each brain region to one emerging cluster only. These modeling approaches are also restricted in exploring new ways to determine the practical and empirical relevance of certain brain regions, such as by linking them to key characteristics of the daily social environment.

For these reasons, autoencoder neural network algorithms are a particularly promising avenue (Danilo Bzdok, Eickenberg, et al., 2015; G. E. Hinton & Salakhutdinov, 2006) to fully appreciate and explicitly model potentially complexe variation across known brain locations that were previously shown to represent the human social brain (Alcalá-López et al., 2018). Hidden subnetwork representations were directly learned from the brain-imaging data themselves by translating autoencoder network solutions from the deep learning community. This artificial neural network approach for pattern discovery inherently yielded empirical validity by gauging the achieved information compression from the variation of the social brain. To show practical relevance, we then probed the social-brain-derived candidate subnetworks by testing their predictive value across a repertoire of diverse human social traits.

## Material and Methods

### Human population data resource

The UK Biobank is a prospective epidemiological resource that provides rich information including brain-imaging, genetics, and multiple biological and lifestyle measurements. We used the brain-imaging data from the 10,000 participant UKB release (see Supplementary Table 1 for demographic information), since this sample was homogeneously recruited at the same assessment centre. We used high-resolution T1-weighted structural magnetic resonance images (MRI), as these measurements can be used to capture whole-brain grey matter morphology (Miller et al., 2016). These brain scans were submitted to preprocessing and quality-control workflows from Alfaro-Almagro and colleagues, FMRIB, University of Oxford, UK (2018). Use of this uniform preprocessing pipeline increases the comparability of our findings to other and future UKB studies. Moreover, we examined several key markers of social lifestyle (Table 1, Supplementary Fig. 1).

**Table 1:** Social lifestyle markers

| **UKBB-ID** | **Social Lifestyle Marker** | **Group 1** | **Group 2** | **Group 3** | **Group 4** |
| --- | --- | --- | --- | --- | --- |
| 2020 | Loneliness | Women feeling lonely | Women not feeling lonely | Men feeling lonely | Men not feeling lonely |
| 4570 | Friendship Satisfaction | Women with low friendship satisfaction | Women with high friendship satisfaction | Men with low friendship satisfaction | Men with high friendship satisfaction |
| 709 | Living Alone | Women living alone | Women living with others | Men living alone | Men living with others |
| 2149 | Number of Lifetime Sexual Partners | Women with one sexual partner | Women with several sexual partners | Men with one sexual partner | Men with several sexual partners |
| 22617 | Job | Women without a social job | Women with a social job | Men without a social job | Men with a social job |
| 2110 | Social Support | Women with low social support | Women with high social support | Men with low social support | Men with high social support |

### Preprocessing of structural brain-imaging data

Structural MRI brain scans (T1-weighted 3D MPRAGE sequence at 1 mm isotropic resolution) were pre-processed using gradient distortion correction, field of view reduction using the Brain Extraction Tool (Smith, 2002) and FLIRT (M. Jenkinson & Smith, 2001)(Mark Jenkinson et al., 2002), as well as non-linear registration to MNI152 standard space at 1 mm resolution using FNIRT (Andersson, J. L., Jenkinson, M., & Smith, S., 2007). To avoid unnecessary interpolation, all image transformations were estimated, combined and applied by a single interpolation step. Tissue-type segmentation into cerebrospinal fluid, grey matter and white matter was applied using FAST (FMRIB’s Automated Segmentation Tool (Zhang et al., 2001) to generate full bias-field-corrected images. SIENAX (Smith et al., 2002), in turn, was used to derive volumetric measures normalized for head sizes. The ensuing adjusted volume measurements represented the amount of grey matter corrected for individual brain sizes.

### Social brain atlas definition

Our study built on a current best-estimate of social brain topography in humans, which only recently became available (Alcalá-López et al., 2018). This topographical atlas was derived by a quantitative large-scale integration of functional MRI findings from 3,972 task experiments involving thousands of individuals. 36 regions of interest were thus previously identified (Supplementary Table 2). These 36 already-established locations were also reported to be connectionally and functionally segregated into four network clusters (Alcalá-López et al., 2018), Fig. 4): i) a visual-sensory cluster (fusiform gyrus, posterior superior temporal sulcus, MT/V5), ii) a limbic cluster (amygdala, ventromedial prefrontal cortex, rostral anterior cingulate cortex, hippocampus, nucleus accumbens), iii) an intermediate cluster (inferior frontal gyrus, anterior insula, anterior mid-cingulate cortex, cerebellum, supplementary motor area, supramarginal gyrus), and iv) a higher-associative cortical cluster (dorsomedial prefrontal cortex, frontal pole, posterior mid-cingulate cortex, posterior cingulate cortex, precuneus, temporo-parietal junction, middle-temporal gyrus, temporal pole).

Our pattern-learning pipelines were thus anatomically guided by brain volume extraction for the 36 consensus brain regions of interest (each associated with one of the four previously established functional clusters in the social brain). In this way, neurobiologically interpretable measures of grey matter volume were obtained in previously established brain locations from the ~10,000 participants (Kernbach et al., 2018; Kiesow et al., 2020 in press; Miller et al., 2016). These values were obtained by summarizing whole-brain structural MRI maps based on the topographical compartments of the social brain. We applied a smoothing kernel of 5mm FWHM to the participants’ structural brain maps to homogenize local neuroanatomical features (Frangou et al., 2004). Grey matter volume information (cf. above) was averaged in spheres of 5mm diameter around the consensus location from the previously established social brain atlas (Alcalá-López et al., 2018), averaging the preprocessed, tissue-segmented, and brain-size-adjusted MRI signals (cf. above) across the voxels belonging to a given target region (Kiesow et al., 2020 in press). This procedure yielded a single representative volume measure for each constituent element of our social brain atlas. Note that using spheres of 2.5mm or 7.5mm diameter yielded virtually identical results and led to the same conclusions.

This approach yielded 36 neurobiologically meaningful volume measures for each UKB participant. Each of these social brain features was z-scored across participants by centering to zero mean and scaling the variance to one. These measures of regional brain volume in social brain networks served as the basis for all subsequent analysis steps. Full information on the social brain locations that provided the basis for this study are available online for transparency and reuse at the data-sharing platform NeuroVault (http://neurovault.org/collections/2462/).

### Neural network algorithms to discover subnetworks hidden in social brain variation

To seize the opportunity to provide a richer picture of potential subnetworks underlying variation across the social brain atlas, we leveraged artificial autoencoder neural networks (Fig. 1). This family of deep learning algorithms can naturally extend to architectures with multiple latent layers of consecutive non-linear processing (Goodfellow et al., 2016; G. E. Hinton & Salakhutdinov, 2006). These algorithms were deployed to extract spatially distributed patterns dormant in the structural MRI data. The representation learning approach directly addressed the question of how the morphological variation across regions of the entire social brain can be re-expressed in a limited set of elementary network representations. This modeling goal was satisfied by imposing a projection of rich input data to a lower-rank bottleneck (Fig. 1) to automatically derive a useful compression of information from structural brain variation into a collection of atomic network patterns (Danilo Bzdok, Eickenberg, et al., 2015; G. E. Hinton & Salakhutdinov, 2006).

**Figure 1.**
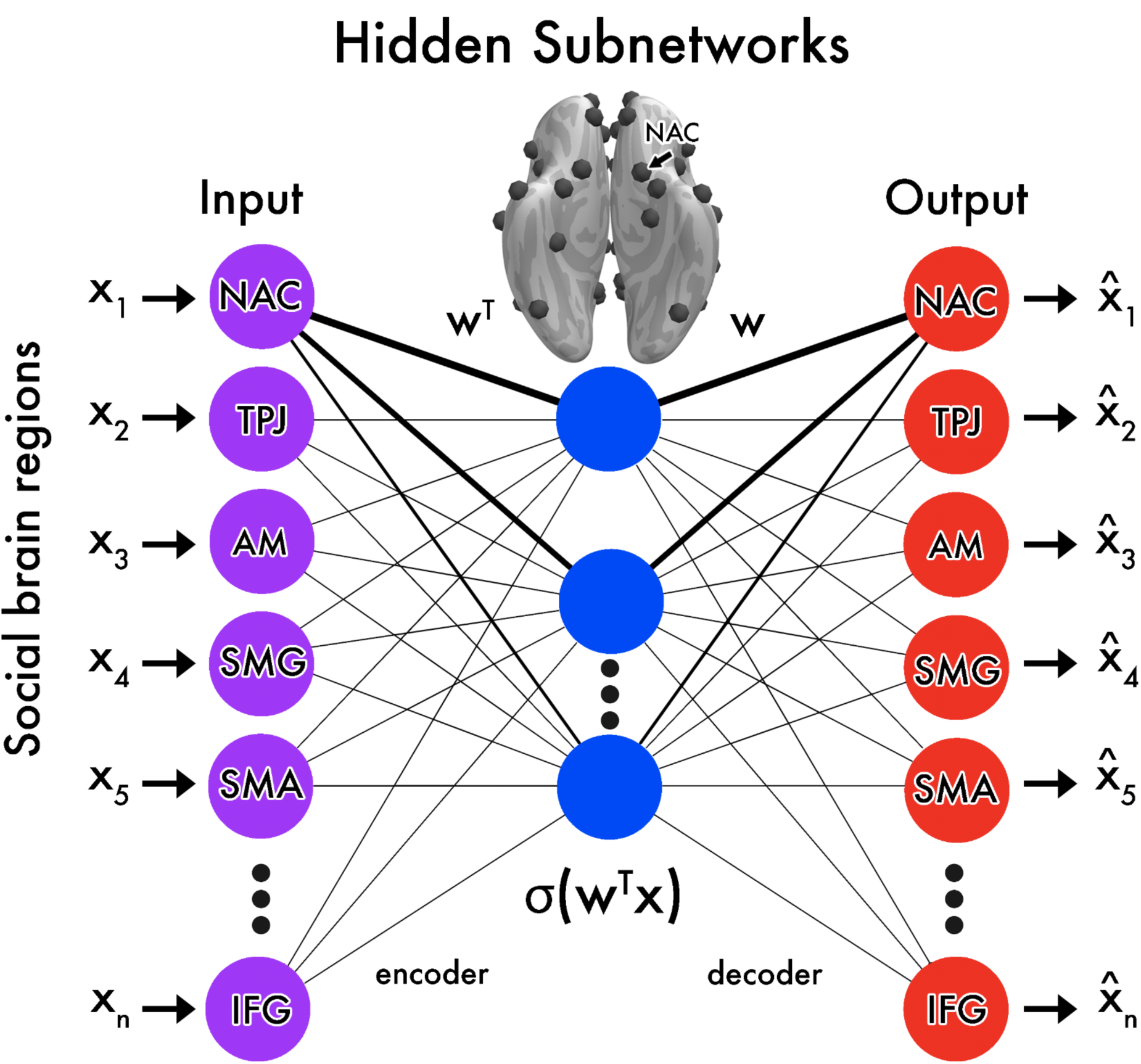
Schematic on how autoencoder neural networks learn to decompose the social brain into structural co-dependency patterns. We designed autoencoder learning algorithms to seek general principles of how regional volume varies cohesively across the social brain atlas (Danilo Bzdok, Eickenberg, et al., 2015; G. E. Hinton & Salakhutdinov, 2006). We assumed that the set of social brain region volumes vary jointly in the broader human population. We therefore explored which of the 36 atlas regions are the core social regions that provide the most information about cohesive volume variation, in an effort to deconvolve the hidden subsystems of structural co-variation in the human social brain. The encoder network (*left*) estimated parameter weights that define the essential structural links from the original region volumes to a re-expression in the learned embedding representation. The so-called bottleneck (*center*) represented the embedding expressions that pool from volume distributions across regions in each unit in the hidden layer. Each hidden unit arises from an adaptive combination of its input links Σ_*j*_ *w*_*j*_ *x*_*i*_ and activation function *σ*(⋅). The decoder network (*right*) estimated parameter weights that learn to use the volume distribution embeddings from the hidden layer to restore each participant’s region volume distribution. Minimizing the discrepancy (reconstruction error) between originally measured and recovered region volumes is the optimization goal that drives the search for artificial neural network solutions. For the purpose of illustration, the nucleus accumbens (NAC) is depicted with its strength of representation (*thickness of links*) in the different emerging social subnetworks (*blue*). As such, the artificial neural network architectures were trained to learn >1,000 parameter weights by asking: ‘Which coherent interregional representations are most instrumental to dis-assemble and re-assemble social brain structure?’

The “encode-decode” modeling scheme yielded one spatially distributed volumetric pattern for each extracted dimension in the bottleneck latent space (Fig. 1). Each of the distributed volumetric patterns encapsulated one hidden subnetwork that quantitatively delineated coherent interregional dependencies across the entire social brain atlas. As such, using one, or up to all extracted hidden subnetworks, the autoencoder could rebuild the regional brain structure that constitutes the human social brain as best as possible. If successful, this modeling agenda can unlock evidence for the subnetworks’ empirically tested ability to parsimoniously recapitulate the brain information wedded into the entire social brain atlas. These artificial neural network algorithms provide an attractive solution for the goal of a comprehensive exploration of hidden sources of variation that collectively comprise the social brain atlas.

Autoencoder learning architectures can be automatically optimized to improve the fidelity of the constituent subnetworks that together, combine to the collapsed measures of social brain volumes that were actually captured using MRI. The optimization objective was based on the original participant volume expressions by means of searching through a vast space of candidate hidden subnetwork patterns to converge on an optimal representational solution. This model family is naturally scalable because these pattern-learning algorithms are well-known to abstract across several classical methods for dimensionality reduction (Danilo Bzdok, Eickenberg, et al., 2015; Goodfellow et al., 2016). In line with the primary goal of our study, the elected modeling framework allowed for each location of the social brain atlas to exhibit a different relevance in different subnetworks. We hypothesized that spatially overlapping subnetworks are critical to making progress towards a faithful representation of brain compartments closely linked to social-affective processing capacities. Our study hence endorses the assumptions that a single target region has a certain association strength with several distinct neurocognitive processes, which accommodates the possibility of mixed membership with continuous degrees of spatial overlap.

To guard against overfitting during model building, we carried out a rigorous cross-validation scheme (Danilo Bzdok, 2017; Hastie et al., 2001). In five (outer) folds of data splitting, structural brain scans from 9,933 participants were randomly divided into a training set (total n = 4,966, 2,575 females, mean age = 55.41 years, SD ± 7.54) and a test set (total n = 4,967, 2,629 females, mean age = 55.27 years, SD ± 7.48). In 10 nested (inner) folds of random data splitting, we used 90% of the training set for model parameter estimation, while 10% of the training set were used for model hyperparameter tuning and model architecture selection. In particular, we charted several architectures of autoencoder neural networks (Table 2) on the volume data centered on the 36 social brain regions. To learn hidden network representations from structural brain scans by means of different autoencoder architectures, we used the RMSprop optimizer (G. Hinton et al., 2012) and a learning rate of 1e-3 based on a grid search of the hyperparameters (see Supplementary Table 3 for details). We probed autoencoder architectures that differed in the number of latent processing layers (i.e., 6, 4, 1), linear versus nonlinear activation functions (identity function versus Relu operation at “neuron” processing units), tied versus non-tied weights and different penalty terms exerting regularization on the weight matrix of the latent layers inside of the layers (l1, l2 and cross-covariance regularization constraints (Cheung et al., 2014)). The process of building hyperparameter-optimized instances of these different artificial neural networks was exclusively performed on the training set (cf. above). In a subsequent step, we evaluated the autoencoder-based information compression performance on unseen participants from the independent test set.

**Table 2.** Examined types of autoencoder neural networks

| **Autoencoder architecture** | **Unit activation function** | **Regularization constraints on parameters** | **Number of latent layers** | **Units per latent layer** | **Tied weights** |
| --- | --- | --- | --- | --- | --- |
| *baseline* | identity | None | 1 | 15 | No |
| *baseline + l1* | identity | l1 penalty | 1 | 15 | No |
| *baseline + l2* | identity | l2 penalty | 1 | 15 | No |
| *baseline + covariance* | identity | covariance | 1 | 15 | No |
| *tied non-linear* | Relu | None | 1 | 15 | Yes |
| *3-layer non-linear* | Relu | None | 3 | 25-15-25 | No |
| *5-layer non-linear* | Relu | None | 5 | 25-20-15-20-25 | No |

### Prediction of social markers based on participants’ subnetwork expressions

We subsequently examined the predictive role of the discovered social subnetworks based on their variation in our population sample. For this purpose, we tested the subnetwork generalizability for several markers of everyday social life (Table 1). For this supervised arm of our analysis workflow, we used the identical nested cross validation procedure (cf. above). That is, inside each of the five outer folds, the particular training set participants were further subdivided into ten splits for the purpose of model selection and model hyperparameter tuning (cf. above). The estimated candidate models were compared against each other on the independent (inner) data splits. This approach allowed identification of the model instance with the best hyperparameter configurations, which was based on the highest achieved relative predictive performance. The built hyperparameter-optimized models were then assessed for their absolute predictive performance on never-seen participant data from the test set (outer loop). To obtain an accurate estimate of the expected performance of the model, the fold-wise model performances were subsequently averaged (i.e., across five separate test accuracies) to a single cross-validated prediction performance, which we expect to hold in other independent or future datasets (Hastie et al., 2009; Pereira et al., 2009).

For the supervised prediction of social lifestyle traits, we charted two classes of predictive algorithms that are complementary in representational capacity and thus theoretically achievable prediction power (Danilo Bzdok & Yeo, 2017). As a widely used classifier with linear capacity, we opted for Tikhonov-regularized regression with logistic loss function. The key hyperparameter of this pattern-learning classifier was the coefficient for the l2 penalty term, which we searched in a grid ranging from −3 to +3 in seven logarithmically spaced steps. As a commonly employed classifier with a considerably higher capacity to detect and exploit complex predictive patterns, we opted for random forest algorithms (Breiman, 2001). For hyperparameter search, we tuned the maximum depth (2 or 6), the minimal split of samples (2 or 6) and the minimum samples of leaves (2 or 6). We noticed that fitting 100 decision trees showed saturation in prediction accuracy based on the out-of-bag estimates on training data from unseen UKB participants by a given decision tree (Hastie et al., 2001). Our rationale was to test for the existence of exploitable non-linear effects in our brain imaging data for predicting social traits. This consideration informed our decision on whether to commit to a high-capacity predictive algorithm, or to resort to a linear predictive algorithm is sufficient for our predictive characterization of the identified subnetworks.

We performed prediction of interindividual differences for a given social trait based on the autoencoder-derived latent factor projections (cf. above) of social brain volume measures. To ensure balanced groups, the UKB participants were split into more social versus less social lifestyles. Each examined social marker was ensured to have binary encoding (median-split as appropriate) into more social versus less social categories. Our approach also acknowledged the wideranging sex differentiation of social traits in the human brain that is receiving increasing empirical support from neuroimaging studies (Kiesow et al., 2020 in press; Tannenbaum et al., 2019). As such, the prediction goal, we further split the participants according to sex, which yielded four groups for classification: 1) more social males, 2) less social males, 3) more social females and 4) less social females. Hence, for each particular index of social richness, our classifiers solved a four-class prediction problem. Moreover, we incorporated age differences into the analysis pipeline by using participant age as an input source of interindividual variation in all predictive models. To enable comparable handling of the multi-class classification problem with both l2-penalized logistic loss and random forest estimators, we used both prediction algorithms in the widely used one-versus-rest scheme (Hastie et al., 2001). In doing so, we obtained parameter weights that indicated the predictive role or contribution for each latent autoencoder embedding of social brain morphology for successfully discriminating UKB participants who live in a more versus less rich social environment.

### Scientific computing implementation

All computations and visualizations were performed in the Python programming language. For the unsupervised arm of the analysis workflow, we used Keras (Chollet & Others, 2015) to create and train the different types of autoencoders, while the predictive algorithms were used as implemented by scikit-learn (Pedregosa et al., 2012). To process and visualize the structural MRI data, we used nilearn (Abraham et al., 2014) and its Pysurfer (https://pysurfer.github.io/) interface. We created all additional figures with Matplotlib (Hunter, 2007), Seaborn (https://seaborn.pydata.org/) and Bokeh (Bokeh Development Team, 2019). Pandas (McKinney, 2010) was used for data slicing and dicing.

## Results

### Neural network algorithms learn coherent subnetworks from social brain variation

We distilled hidden subnetwork representations from structural variation across the social brain atlas in ~10,000 UK Biobank participants. This goal was achieved by charting several artificial neural networks that implement autoencoder variants. For a given algorithm architecture, the information compression performance was computed by invoking back-projection from each participant’s specific subnetwork embedding expressions to recover volume estimates for all 36 social brain atlas regions (Bzdok et al. 2015). That is, we computed the difference between the actual volumes of each social brain region as measured in each participant and the volumes reconstructed from the participant-wise hidden subnetwork expressions as a metric of parsimony of the derived candidate representations.

The probed autoencoder neural networks varied in key properties including the depth of the consecutive processing layers, intricacy of modeled intervariable relationships, and different regularization constraints on model parameter estimation (Table 2). Among the deep non-linear autoencoder architectures with Relu activation function, the six-layered autoencoder achieved a high explained variance (EV) of 0.78 (SD < 0.01, measured as mean absolute error). The baseline autoencoder architecture with identity unit activation function and one latent processing layer and without regularization constraints was the simplest architecture, and also achieved an explained variance of 0.78 (SD < 0.01 across data splits) (Fig. 2). Hence, the information compression performance of the baseline autoencoder in learning a parsimonious representation of the original social brain regions was not outperformed by other probed autoencoder architectures based on the explained variance (i.e., mean absolute error) or stability (i.e., standard deviation over different data splits) (Fig. 2). As deeper non-linear neural network algorithms did not yield statistically defensible performance improvements on our structural brain data, we focused on baseline autoencoder neural networks with one latent layer for all subsequent analyses.

**Figure 2.**
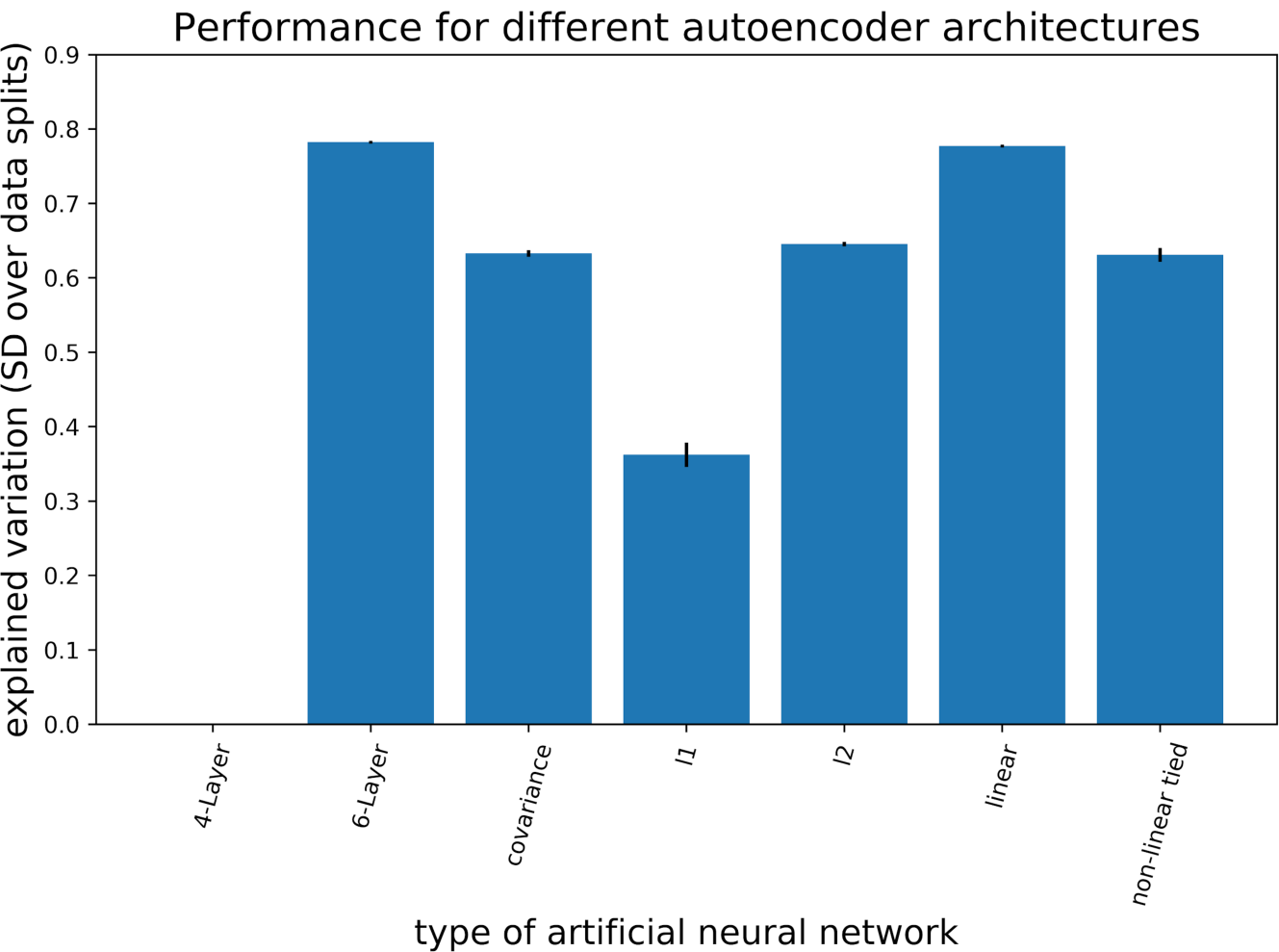
Model performance for different autoencoder neural networks from deep learning. After forming a given autoencoder model based on the training participants, all models’ explained variance performances (i.e., mean absolute error) were evaluated on independent test participants. The bar plot shows that the baseline autoencoder (linear processing layers with identity activation functions) performs at least as well as any of the other probed, more sophisticated neural network algorithms. The mean explained variance is indicated as the heights of the blue bars. The *error bars* display the stability of performance as measured by standard deviation (SD) across data splits. The more complex 6-layer autoencoder architecture was not able to outperform the baseline autoencoder solution in a convincing way. This observation probably witnesses the risk of overfitting in more complex autoencoders. Hence, for our study, there is no substantial advantage of deep or nonlinear autoencoders. Furthermore, different penalties did not improve the model. In terms of both model performance (i.e., explained variance) and stability (i.e., SD), the baseline autoencoder was at least as successful at extracting structural dependence patterns in the social brain as any other examined autoencoder architecture.

By means of the baseline autoencoder neural networks with identity unit transformations (instead of Relu activation function), we examined the effect of different types of regularization constraints on compression performance. For the purpose of increasing sparsity, we encouraged exactly-zero parameter values during model estimation, corresponding to region relevances, by imposing l1 regularization (EV = 0.36, SD ± 0.02). To instead constrain the estimation of model parameters towards smaller absolute values, we imposed l2 regularization, which yielded better model performance (EV = 0.65, SD < 0.01). Finally, constraining the network pattern discovery to discourage mutual correlation between the emerging subnetworks using a covariance penalty term yielded performance (EV = 0.63, SD < 0.01), which ranked in-between that of l1 and l2 penalized neural network algorithms. Hence, imposing different types of regularization constraints on the baseline autoencoder did not outperform the information compression performance of the overall social brain morphology in unseen data, and also led to similar solutions of hidden subnetwork representations.

In a series of similarity tests, we additionally assessed the robustness of our candidate subnetwork solutions for the social brain. Within each of the subnetworks, we compared the relevance patterns of the social brain regions to corresponding hidden representations emerged from the other autoencoder variants with different regularization constraints. The robustness of all assigned region relevances to the derived hidden representations was suggested by subnetwork-wise Pearsons’ correlations across the 36 region relevances that were learned by different autoencoder architectures (Sup. Fig. 2, Sup. Fig 3 A-C). The Pearson correlation coefficients averaged over four (Sup. Fig. 3A-C) different autoencoder neural networks with identity function was rho = 0.97 (SD ± 0.05). In particular, the most deviant architecture among these autoencoders was with covariance penalty loss, which showed a mean Pearson correlation of rho = 0.92 (SD ± 0.03). Additionally, a similar Pearson correlation of rho = 0.98 (SD ± 0.01) was obtained when comparing a given type of identity-function autoencoder neural network obtained from the training set with the corresponding architecture learned on the independent test set (Sup. Fig. 3D). Together, these confirmatory analyses ascertained that the region relevances derived by the baseline autoencoder were stable over several neural network architectures. All subsequent analysis steps hence placed focus on the baseline autoencoder with an unconstrained parameter estimation (i.e., without penalty for parameter regularization).

After considering the achieved explained variance and confirming robustness of different types of baseline autoencoders, we directed attention to how much information each separate subnetwork carries about variation in the whole social brain structure. In this series of analyses based on the baseline autoencoder, subnetworks 7, 9, and 15 emerged as most dominant. An elbow-shaped pattern after the third top subnetwork (subnetwork 15) showed a drop in information compression performance (Fig. 3, left panel). Put differently, subnetworks 7, 9 and 15 were highlighted as the three top hidden subnetworks because these specific hidden representations showed the highest importance for encapsulating variation in the complete social brain from only a few hidden structural patterns. It is an important quality of this analytical approach that each social brain region can potentially contribute to multiple subnetworks. This property allowed the set of hidden subnetworks to model several overlapping sources of population variation at the same time (Fig. 4).

**Figure 3.**
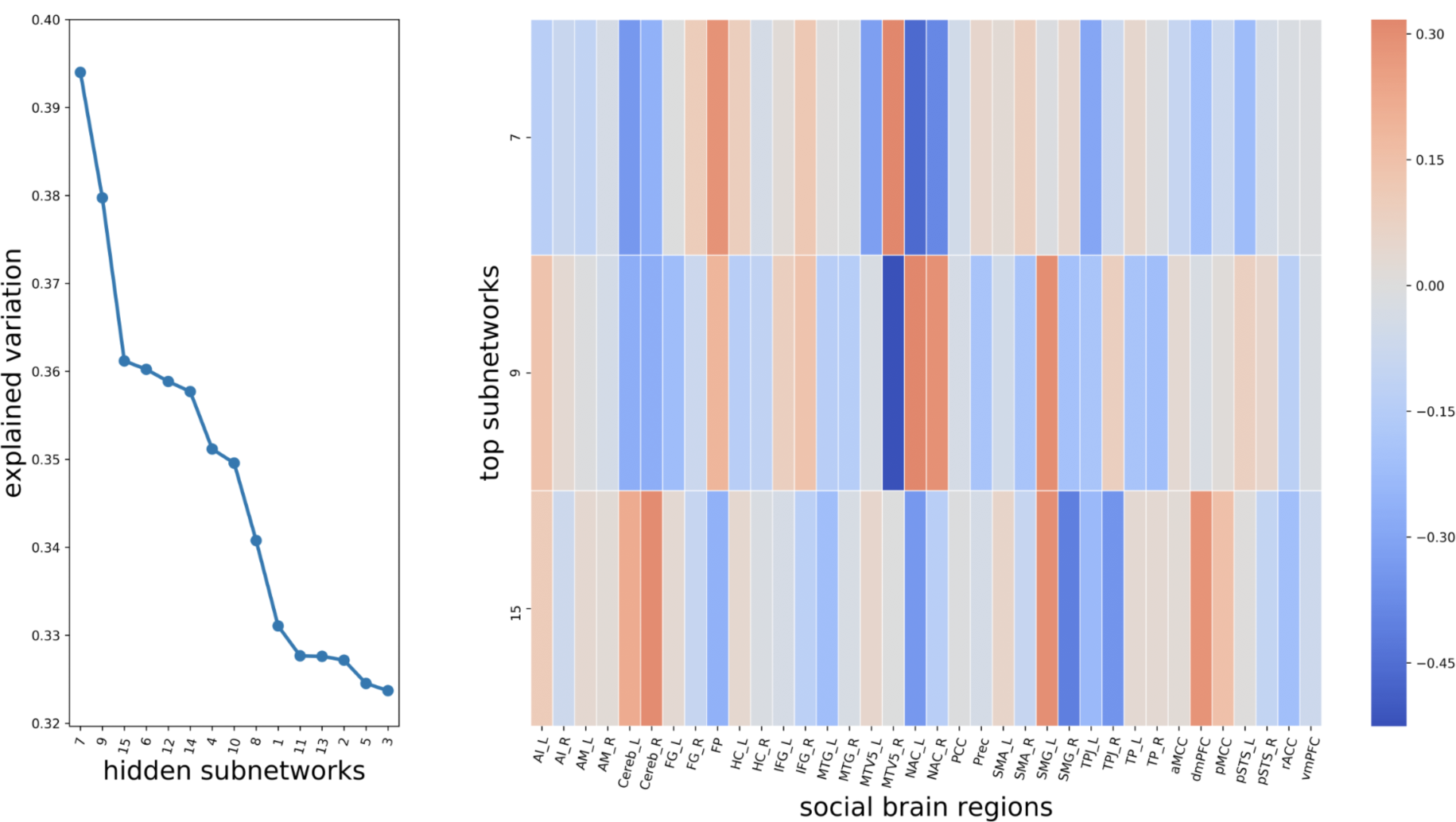
Most explanatory hidden subnetwork representations learned by the autoencoder neural network. *Left*: We quantify the volume effects of each particular social subnetwork to volume variation in the whole social brain atlas. One-after-one, each identified hidden subnetworks (*x axis*) served separately to reconstruct the originally measured volume of 36 social brain regions from each individual. The reconstruction performance of social brain volumes is assessed based on mean absolute error (*y axis*) for each hidden subnetwork. A perfect encapsulation of the complete social brain volume would yield an explained variance of 1, while incomplete recovery would yield numbers lower than 1. This explained variance metric quantified to what extent the directly measured social brain volumes could be restored by one of the learned, not directly measurable subnetwork patterns. *Right*: The 36 social brain regions (*x axis*) contribute differently to the three dominant subnetworks (*y axis*). The relevance of each particular brain region to the dominant subnetworks is represented by the color for a respective combination of regions in a given subnetwork. A positive (negative) relevance towards regional volume is shown in *red* (*blue*) tones. The directionality of the volume effects shows which regions have opposite effects in explaining social brain variation. For instance, on the one hand, the nucleus accumbens is highlighted in the leading three subnetworks 7, 9 and 15, which exemplifies the possibility of overlapping volume variation. On the other hand, of the top three subnetworks, the dorsomedial prefrontal cortex are highlighted in subnetwork 15. For abbreviations and details on social brain regions see Supplementary Table 2.

**Figure 4.**
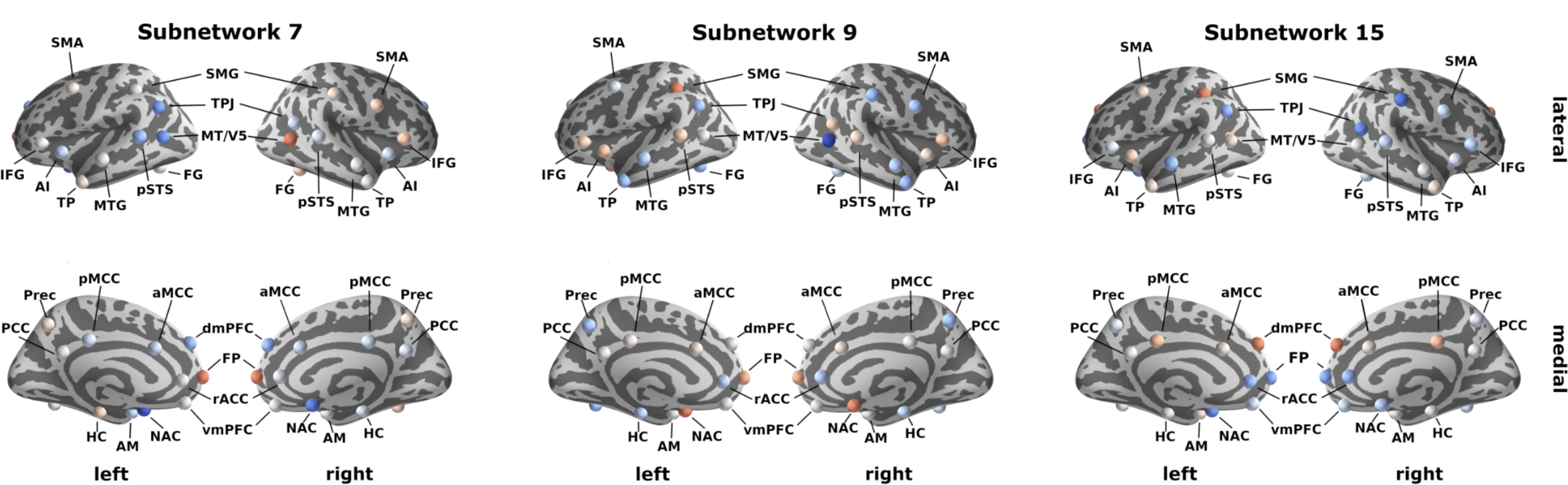
Top three hidden subnetworks that underpin structural dependence patterns in social brain differences. Depicts the uncovered hidden representations that were learned by the autoencoder neural network, with the allocated relevance of each region from the social brain atlas (cf. Fig. 3). *Red* (*blue*) colors represent the extent of positive (negative) relevances. The strongest three hidden subnetworks 7, 9, and 15 represented meaningful structural inter-dependencies that highlighted key regions including the TPJ, medial PFC and NAC. For abbreviations and details on social brain regions see Supplementary Table 2. Color coding according to Figure 3.

Consequently, the particular set of region volume effects in a specific subnetwork should be interpreted in light of the relevances inside of the other concomitant subnetworks (Supplementary Fig. 4), which compose the overall human social brain. In particular, the nucleus accumbens (NAC) contributed strongly to all three dominant subnetworks 7, 9, and 15 (Fig. 3, right panel). The bilateral MT/V5 yielded similar region relevances for subnetwork 7. Additionally, the right MT/V5 contributed strongly to subnetwork 9. Moreover, subnetwork 7 allocated region relevance to the temporo-parietal junction (TPJ) and frontal pole (FP), while subnetwork 9 also highlighted the relevance of the bilateral supramarginal gyrus (SMG). Furthermore, the bilateral SMG, bilateral TPJ and dorsomedial prefrontal cortex (dmPFC) also substantially contributed to subnetwork 15.

To functionally annotate the learned hidden subnetworks (cf. methods), we then performed a descriptive characterization in context to the previously established functional clusters in the social brain atlas (Alcalá-López et al., 2018). This previous study grouped the 36 constituent regions into four hierarchically differentiated functional circuits: the visual-sensory, intermediate, limbic, and higher-associative clusters. We computed the aggregated (absolute) relevances of all social brain regions inside of each previously defined cluster for the present social subnetwork representations (Figs. 4 and 5). For subnetwork 7, the highest aggregate relevances were found for the visual-sensory cluster (0.17 on z-scale, cf. methods) and limbic cluster (0.14). For subnetwork 9, similar aggregate relevances were apparent across all four hierarchical clusters, however the visual-sensory (0.16) and intermediate clusters (0.16) yielded the highest identical aggregate relevances. For subnetwork 15, the intermediate cluster yielded the highest aggregate relevance (0.16) followed by the higher-associative cluster (0.15).

**Figure 5.**
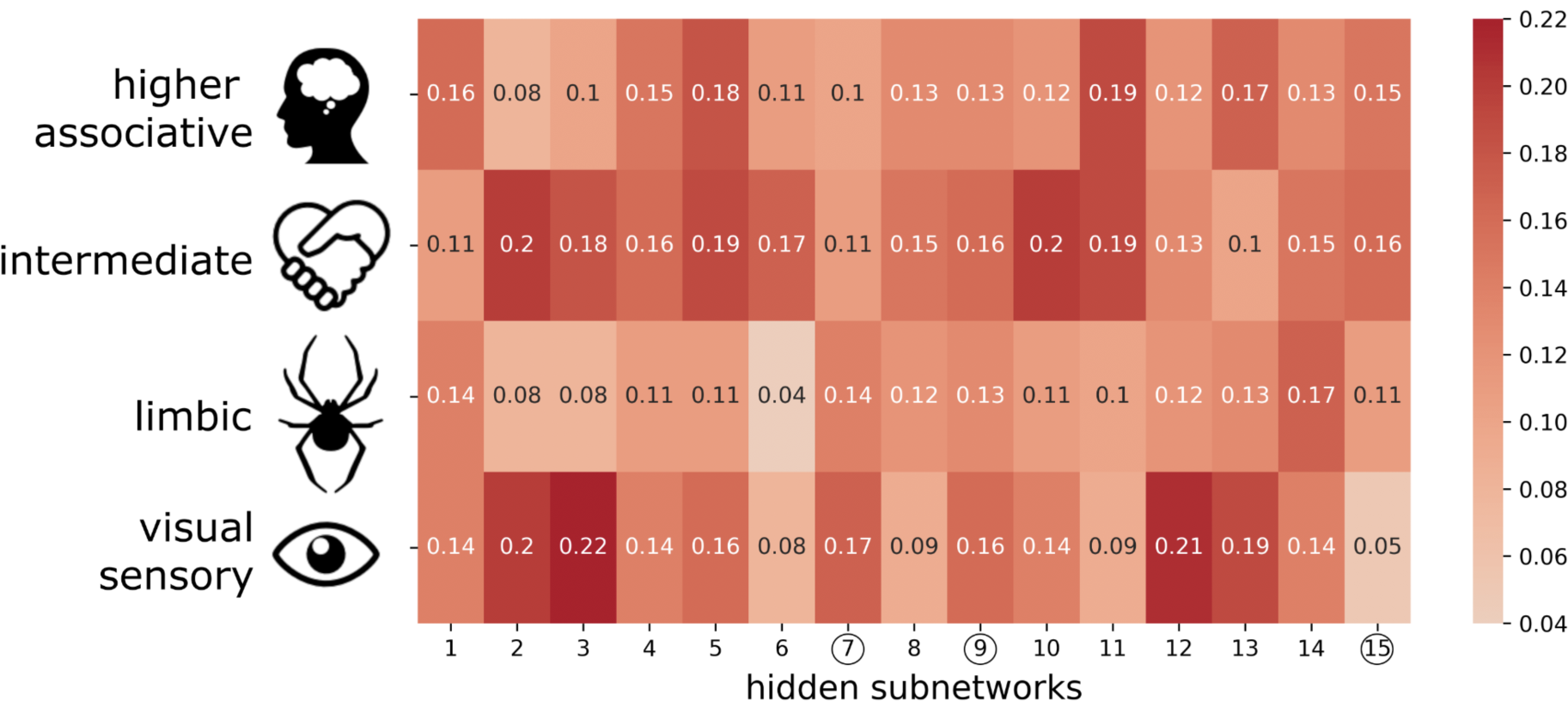
Functional and hierarchical annotation of the learned hidden social subnetworks. To provide additional functional profiling, we related the derived subnetworks to previously reported clusters, which correspond to four decreasing hierarchical levels of social brain circuits: higher-associative, intermediate, limbic and visual-sensory systems (Alcalá-López et al., 2018). The *red* color indicates the cluster-per-cluster relevance that are parsed for the social subnetworks derived in the present study (*x axis*). This metric was calculated as the average (absolute) relevance of all brain regions part of a consensus cluster (colorbar on z-scale). For instance, in subnetwork 15, aggregated relevance is distributed relatively evenly between higher-level and intermediate functional systems. In contrast, subnetworks 3, 7, 9, 12 and 13 show stronger relations to the lower visual-sensory functional cluster. These alternative summaries of our results validate the previously investigated influence of these consensus clusters. These post-hoc exploratory results also highlight that we are able to build upon this existing knowledge with overlapping representations of different underlying subnetworks.

To summarize the unsupervised analyses on subnetwork discovery, if we only had access to each participant’s volume expression from the three most dominant hidden social subnetworks, we could produce a reliable estimate of the complete social brain morphology across UKB participants. That is, the regions assigned with strongest volume effects in these dominant three subnetworks are at the core of the interregional structural dependencies that combine to empirical measures of social brain variation.

### The discovered subnetworks forecast diverse facets of social life

In the supervised arm of our study, we finally sought understanding of the predictive profiles of the learned social subnetwork representations for relevant indicators of social lifestyle. For this purpose, we assessed each participants’ individual combination of subnetwork expressions as a basis for classifying social traits that have an impact on interindividual variation in everyday social interactions (Fig. 6). Across the examined social markers, our predictive pattern-learning algorithm distinguished more versus less sociality in males and females. Individuals who were socially less satisfied, had fewer social interactions or indicated a lower quality for a given social marker were assigned to the less social group. Instead, those more socially satisfied or with more opportunities for social interaction were assigned to the more social group. We used a four-class linear classification approach, where each fitted instance of the pattern-learning algorithm predicted one group (e.g., social females) against the three remaining groups (e.g., non-social female, and social or non-social male) given the participant-specific embeddings of subnetwork expressions.

**Figure 6.**
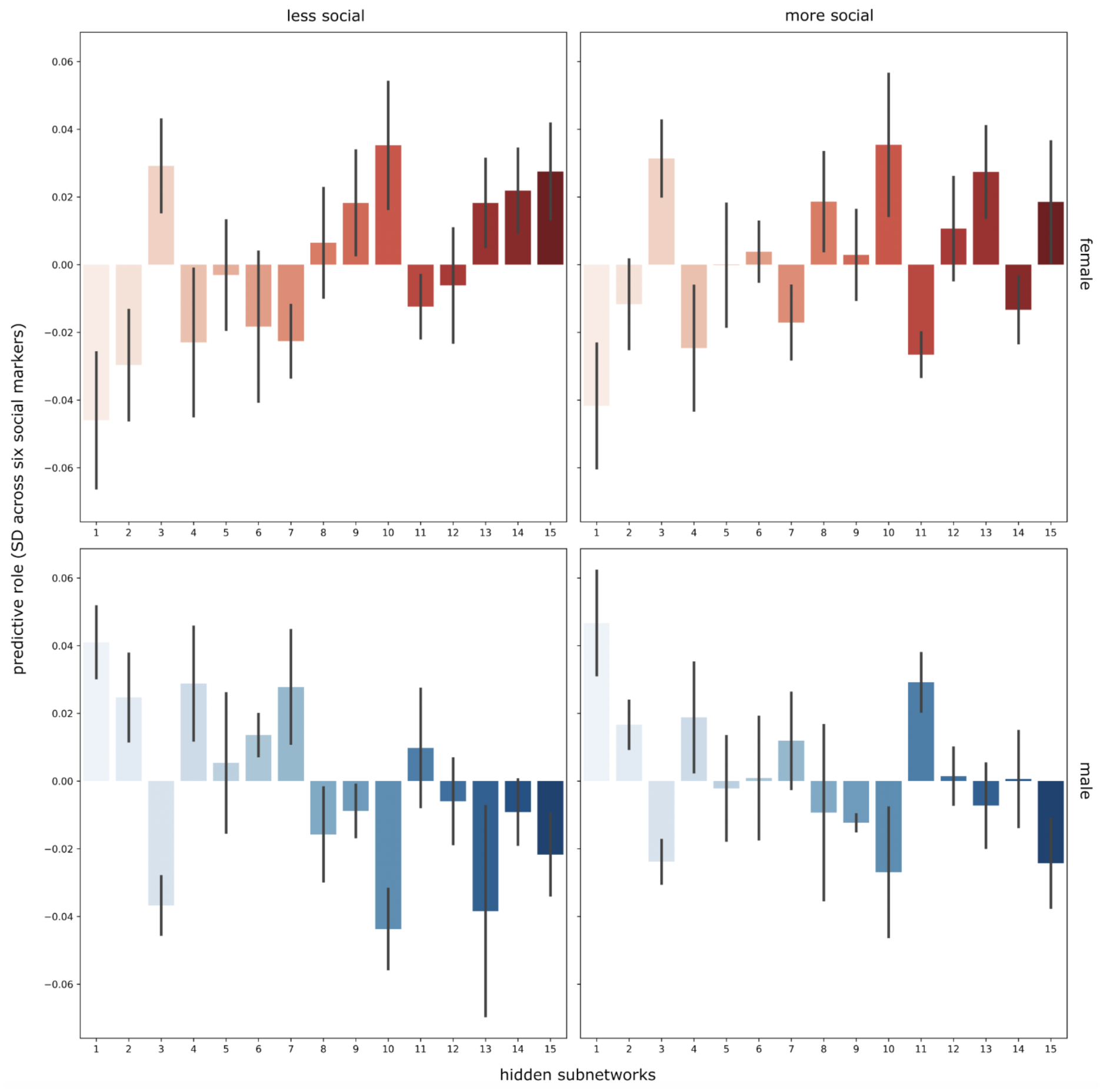
Overall predictive role of the hidden subnetworks for tracking more versus less social exchange. Across all examined social markers, each bar indicates the predictive contribution (*y axis*, units on z-scale) of a given subnetwork (*x axis*) for the degree of sociality in female and male participants. A logistic-loss classification algorithm was trained based on variation in subnetwork expressions across participants to learn predictive patterns for distinguishing the amount of regular social stimulation in men and women (*left*: less social, *right*: more social, *red*: female, *blue*: male). Our analytical approach thus yielded one set of subnetwork weights for each of the four target groups to be classified (*four panels*). The obtained classifier weights are summarized across all social traits (error bars for each subnetwork, SD=standard deviation). The predictive contributions (*y axis*) corresponding to each subnetwork (*x axis*) are shown in each bar. Several subnetworks (e.g., 15, 10, and 1) showed strong predictive contributions across analyses. In general, the directionality of each prediction weight appears to be more tuned to sex, while the relative differences in prediction weights are more tuned to the richness of social environment.

We then tested whether a nonlinear classification algorithm could outperform our simpler linear classifier (cf. methods) by leveraging potentially exceedingly complicated patterns in social brain variation at population scale. To this end, we used random forest algorithms as a higher capacity estimator to assess the out-of-sample prediction performance of participants’ social traits based on the participants’ subnetwork expressions. Virtually identical prediction accuracies in new participants were observed for both logistic-loss classifier (classification accuracy = 0.29, SD ± 0.02 across data splits) and elaborate random forest classifier (classification accuracy = 0.30, SD ± 0.03). Note that both classes of predictive algorithms performed better than the chance level of 0.25 in this four-class scenario. However, given the similarity in out-of-sample performance, its overlapping dispersion, and our goal of direct interpretability of most discriminatory social subnetworks, we embraced the simpler logistic-loss classifier for all subsequent analyses.

Across all examined social traits (Fig. 6), interindividual variation in hidden subnetwork 1 (characterized by high relevance of fusiform gyrus, frontal and dorsomedial prefrontal cortex and posterior mid-cingulate cortex, cf. Sup. Fig. 4) was particularly informative for detecting differences in regular social experience (predictive model weight *w*_*1*_ = 0.04, SD ± 0.02 across social traits). Instead, variation in subnetwork 10 (high relevance of anterior insula, rostral anterior cingulate cortex, and supramarginal gyrus) appeared especially tuned to sex differences based on its predictive contribution to the classifier (*w*_*10*_ = 0.04, SD ± 0.02), rather than showing salient trait-specific patterns (Supplementary Fig. 5). In line with our study goal, we therefore focused attention on the hidden subnetworks with predictive roles for different social markers. These were the trait-discriminatory social subnetworks 3 (*w*_*3*_ = 0.03, SD ± 0.01), subnetwork 4 (*w*_*4*_ = 0.03, SD ± 0.02), subnetwork 13 (*w*_*13*_ = 0.02, SD ± 0.02), subnetwork 15 (*w*_*15*_ = 0.02, SD ± 0.01), subnetwork 11 (*w*_*11*_ = 0.02, SD ± 0.01), subnetwork 2 (*w*_*2*_ = 0.02, SD ± 0.01) and subnetwork 7 (*w*_*7*_ = 0.02, SD ± 0.01). Thus, volume variation of these hidden subnetworks was the most useful for accurately predicting interindividual differences in social exchange.

In addition to predicting participants’ overall degree of sociality, we next zoomed in on the hidden subnetworks that were able to best predict specific markers of social life (Fig. 7). To tell apart whether participants were lonely or not lonely, interindividual variation in hidden subnetwork 13 (high relevance of right temporoparietal junction and left posterior superior temporal sulcus) emerged as most useful (e.g., lonely men: *w* = −0.09, SD ± 0.01, more surrounded men: *w* = 0.00, SD ± 0.01 across data splits), in addition to that of subnetwork 3 (high relevance of right supramarginal gyrus, left temporo-parietal junction, and bilateral posterior superior temporal sulcus) and subnetwork 1 (cf. above). For discriminating participants living alone from participants with richer social interaction at home, subnetworks 10 (cf. above) and 4 (high relevance of left temporoparietal junction, bilateral temporal pole, bilateral cerebellum) yielded the relatively highest predictive role (e.g., subnetwork 4: women living alone: *w* = −0.05, SD ± 0.01, women living with others: *w* = −0.01, SD ± 0.01). Subnetworks 1 (cf. above) and 10 achieved the highest predictive roles for differentiating the social brain morphology of participants with regular exchange with peers for social support (e.g., subnetwork 10: men without social support: *w* = −0.06, SD ± 0.01, men with social support: *w* = −0.03, SD ± 0.00). Both subnetworks 1 and 10 also showed individual predictive roles for high versus low self-reported satisfaction with friendship circles. For disentangling volume patterns in the social brains of participants with more versus less daily social interaction at work, salient predictive contributes were identified for subnetwork 10 and subnetwork 1 (e.g., women without a social job: *w* = −0.03, SD ± 0.00, women with a social job: *w* = −0.06, SD ± 0.00). Interindividual morphological variation in social brain structure for both subnetwork 1 and subnetwork 3 played the biggest predictive role for participants with more monogamous versus more promiscuous romantic relationships (e.g., subnetwork 1: women with one romantic partner: *w* = −0.05, SD ± 0.01, women with more romantic partners: *w* = −0.01, SD ± 0.01).

**Figure 7.**
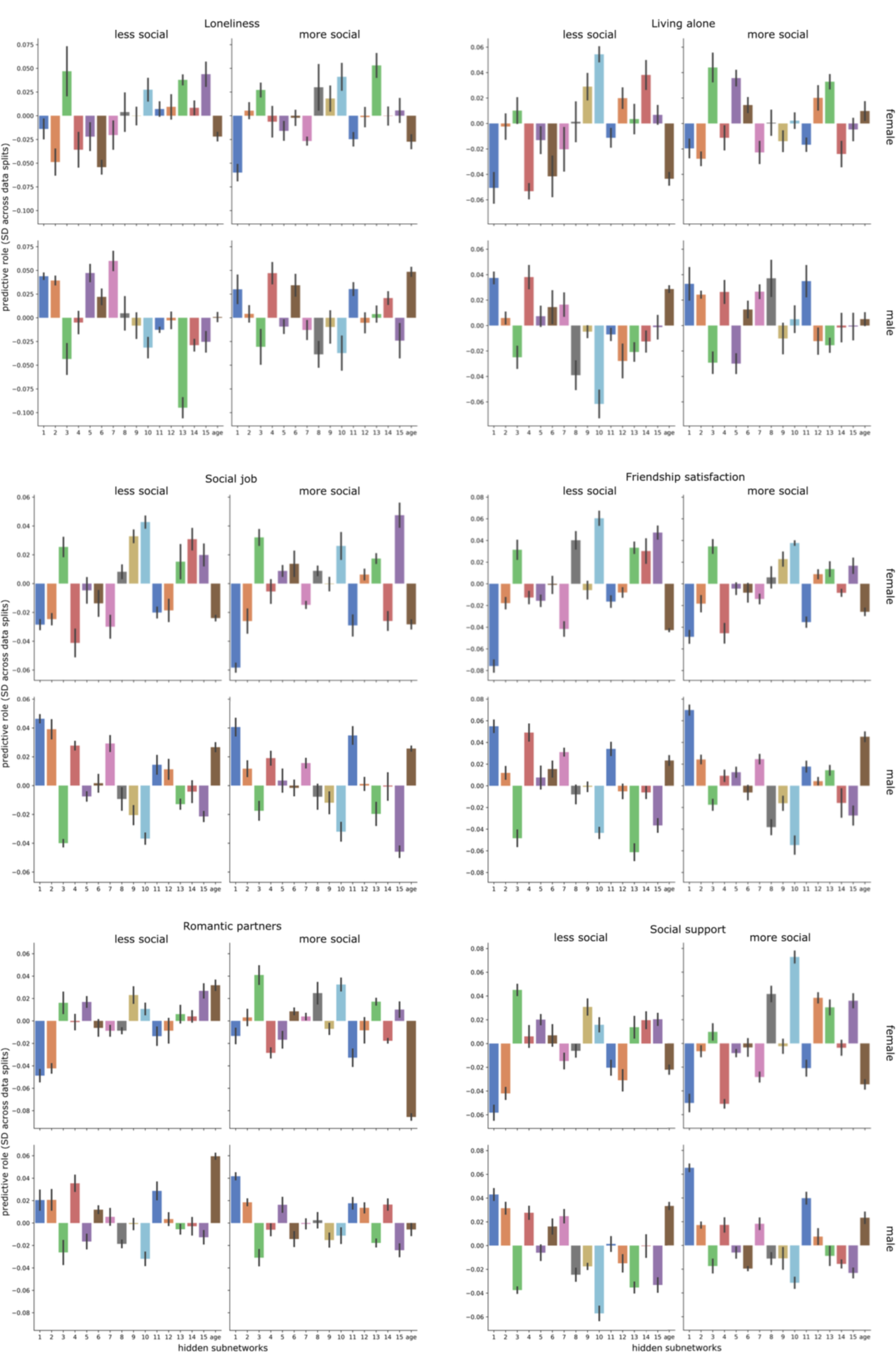
Specific predictive profile of the hidden subnetworks for tracking single social markers. The classification algorithm (cf. Fig. 6) was applied to learn predictive patterns separately for each social marker (*six larger panels*). Each plot depicts the hidden subnetworks (*x axis*) with their predictive contributions (*y axis*, units on z-scale). Each application of the analytical approach yielded one set of subnetwork weights for each of the four target groups to be classified (*four smaller panels*). The obtained classifier weights were summarized for each subnetwork across cross-validation (CV) data splits (*error bars*).

In sum, all revealed hidden social subnetworks showed specific predictive roles comparing between the examined social markers. Notably, the hidden subnetworks 1 and 10 most frequently achieved among the highest predictive roles for specific individual social markers. As such, each source of population variation in social brain structure reliably tracked largely distinct aspects of everyday social interaction in the family, during leisure time and at work.

## Discussion

We set out to uncover elementary building blocks that underpin social brain differences at the population level. The delineated top network representations hidden in the social brain atlas distilled information from sources of population variation and effectively recapitulated the total social brain structure across individuals from the UK Biobank cohort. Specific social brain regions embedded within each subnetwork emerged as especially informative about cohesive dependencies that describe volume variation across the entire social brain. As a common denominator across several extracted subnetworks, the NAC, TPJ, and medial PFC emerged as core network regions in explaining configurations of mutual dependence in social brain morphology. We show how these separable brain representations distinctly predict indicators of everyday social life, such as the subjective experience of loneliness. These signatures of cohesive interregional variations became apparent by algorithmically disassembling and re-assembling structural features of the social brain using autoencoder neural networks.

Many hypothesis-driven social neuroscience studies relied on a set of canonical cognitive concepts for their analysis and interpretation of neural effects. To flank these theory-guided efforts, the present pattern-learning investigation translated algorithmic techniques from the deep learning community. We empowered pattern discovery in the social brain by autoencoder neural networks (Danilo Bzdok, Eickenberg, et al., 2015; G. E. Hinton & Salakhutdinov, 2006). This under-exploited algorithmic technique unlocked insight from uniformly acquired brain scans of the largest brain-imaging cohort recruited from across the United Kingdom.

In our study, the NAC emerged as one of the driving factors of how social brain regions coherently co-vary with each other across thousands of participants, which became apparent in all leading social subnetworks. Traditionally recognized to be implicated in reward-guided decision-making processes, a host of social neuroscience research suggests that the NAC is also one of the core brain regions that are consistently recruited to also support rewarding aspects of social interaction (Behrens et al., 2009). For instance, a functional brain-imaging experiment reported striatal activity in response to both receiving monetary rewards and receiving positive feedback about one's own trustworthiness by unknown others (Izuma et al., 2008). The authors suggested that social approval from others, such as feedback about one’s own personal reputation, may have a common neural basis with nonsocial rewards. In addition, the authors also observed medial prefrontal activity only during the social reward trials, suggesting the mPFC may be specifically involved in the management of one’s own reputation (Izuma et al., 2008). In line with our findings, the NAC and mPFC were flagged as highly relevant in the dominant hidden subnetworks. Consistently, the dominant subnetworks also showed predictive roles for rewarding aspects of social interaction such as friendship satisfaction and having an occupation with frequent social contact.

Indeed, a functional neuro-imaging study assessed neural activity in response to simulating social interactions with friends versus celebrities in an approach-avoidance experiment (Güroğlu et al. 2008). The study showed neural activity responses in the mPFC, NAC, TPJ, posteromedial cortex, SMG, and occipital-temporal junction extending into the MT/V5, specifically when participants interacted with their friends (Güroğlu et al. 2008). The authors (Güroğlu et al. 2008) interpreted that the encounter with close friends may encourage recruitment of interpersonal processes such as empathy, emotion-regulation and reward, all of which may contribute to mental health and positive well-being in the long run. Thus, these findings from functional brain-imaging experiments are in line with our data-led structural findings, and especially highlight the NAC, mPFC, SMG and other co-occurring subnetwork representations that resulted from our social brain decomposition. This set of regions were also placed at the heart of meaningful structural inter-dependencies in our leading hidden subnetworks. Furthermore, our findings revealed these key regions to support prediction of interindividual differences in social lifestyle markers.

Neural activity responses in the NAC are not usually thought to encode differences in intentions of the interaction partner per se (Behrens et al., 2009; D. Bzdok et al., 2011). Instead, perspective-taking processes are typically attributed to a set of higher-level cortical regions with prominent involvement of both the TPJ and mPFC. For instance, a previous structural brain-imaging study identified an association between the ability to read the mind of others through the eyes and grey matter volume in the mPFC, posteromedial cortex and TPJ (Sato et al., 2016). The authors suggest that these social brain regions may contribute to processes necessary to subserve the ability for mental state inference by reading people’s eyes. We extend these previous findings by invigorating the special role of the mPFC, posteromedial cortex, and TPJ in brain circuits related to human social interaction from the present view on the social brain through the lense of subnetworks: The mPFC, posteromedial cortex, and TPJ here explained notable volume effects to major sources of population variation in the social brain, especially in our top subnetworks 15, 7, and 9. In addition, the TPJ was also highlighted as part of subnetwork 13, which we found to help predict loneliness in UK Biobank participants.

In addition to the TPJ as a region critical for realizing high-level social thoughts like perspective taking, a parallel line of research has instead emphasized the mPFC in many other forms of social interaction (Danilo Bzdok et al., 2013; Eickhoff et al., 2016; Schurz et al., 2014). For instance, a series of brain-imaging studies have linked the relationship between several socially responsive regions including the mPFC and indices of interpersonal phenomena (Kiesow et al., 2020 in press; Lewis et al., 2011; Powell et al., 2010), which are reminiscent of our present findings on friendship satisfaction and social support. For instance, a previous structural brain-imaging study mapped grey matter volume in the mPFC, TPJ and STS to intentionality ability and social network size, suggesting these brain regions to be key neuroanatomical correlates for social skills (Lewis et al., 2011). Our current results underscore such findings by showing these social brain nodes to represent major sources of population variation, with overlapping volume effects from several subnetworks. This became apparent in region relevances in the hidden subnetworks 5, 6, 8 and 14, as well as the dominant subnetworks 7 and 15. Additionally, the mPFC and other structurally coupled regions have also been found to be linked to anticipating social feedback. For instance, one functional brain-imaging study reported the mPFC, posteromedial cortex, visual association cortex extending into the MT/V5 and ventral striatum, encompassing the NAC, to show more activity when anticipating positive social feedback from novel peers (Powers et al., 2013). This observation suggests that in concert with the NAC and MT/V5 regions, the mPFC may also play a critical role in navigating salient social encounters (Powers et al., 2013). Thus, our investigation confirms and details the central position of the TPJ, mPFC as key drivers in co-occurring neural substrates that support neurocognitive facets central to social behavior.

Little existing data-driven evidence appears to simultaneously focus on the relevance of the mPFC and TPJ to social cognition, perhaps in part due to the location-by-location logic of most brain-imaging task studies. As one of few exceptions, Schurz and colleagues (2014) conducted a coordinate-based meta analysis of various functional brain-imaging experiments using various psychological paradigms to probe perspective-taking, including social animations, reading the mind in the eyes, and trait judgment tasks. The authors identified foci of meta-analytically derived hotspots of neural response averages that yielded activity convergences situated in the TPJ and mPFC regions. Different from our approach, Schurz and colleague (2014) used pre-existing topographically distinct clusters based on structural connectivity from diffusion weighted brain imaging. Such clusters have strict topographical boundaries that are mutually exclusive, which conveys rigid a-priori assumptions about what to expect in brain-imaging data like MRI scans (Danilo Bzdok et al., 2016). Hence, many previous brain-imaging studies may have ignored possibly overlapping biological phenomena; and joint volume effects of a particular region volume on the complete social brain morphology. On the interpretational level, Schurz and colleagues (2014) suggested the mPFC to play a role in maintaining mental representations of another person’s social and emotional vantage point to create a model of another person's mental life. Our present results allow re-contextualization and provide solid grounding for such localizationist interpretations in mutually overlapping subnetwork representations, which we show to vary in distinct ways at the population level and be differently associated with markers of social richness.

Compared to this previous study, a Bayesian latent factor meta-analysis is more closely aligned with our present analysis tactic. Yeo and colleagues (2015) examined mutually overlapping components of neural activity with a topographical focus on the higher association cortex and its relation to a general battery of task responses. The study answered which of 83 different experimental paradigms, including the n-back test, Stroop test and anti-saccade tasks, exhibit concomitant neural activity changes according to the identified underlying spatially distributed neural activity components. This previous study singled out one functional activity component (component 10), which turned out to be preferentially linked to social cognition. This neural activity component isolated the mPFC, posteromedial cortex, the SMG, and TPJ, all of which were also highlighted in several extracted social brain subnetworks. We complement this previous investigation of general cognitive domains in the higher association cortex by showing coherent structural configurations from a data-driven decomposition of the whole social brain in a larger participant sample, which closely represents the wider UK population.

More broadly, previous cross-modal brain-imaging research has shown that the regions belonging to the human social brain can be hierarchically grouped into a) lower sensory, b) limbic, c) intermediate, and d) higher associative neural systems (Alcalá-López et al., 2018). The described functional compartments were derived under the strict assumption that each social brain region is assigned to only a single group at once. To relax such discrete one-to-one responsibilities, our analyses explicitly quantified the continuous degrees to which a specific subset of social brain regions are relevant in explaining structural variation of multiple subnetworks. Such degrees of multi-to-multi responsibilities therefore allow for each subnetwork to allocate relevance to several of these neural circuits in the social brain. In addition to the TPJ and mPFC, other examples for such regions include the SMG, which has notable relevance in several of our subnetworks. Despite the prevalence of specific brain regions to be relevant in several subnetworks, other subnetworks allocated region relevances more evenly to different functional compartments. For instance, the previously established visual-sensory circuit of the social brain was here most associated with subnetworks 3, 12 and 13. The specificity of such functional annotations is illustrated by the observation that subnetworks 4 and 14 allocated relevance quite evenly between all subsystems. As such, we were not only able to show the prominence of single functional compartments in specific subnetworks, but also an overlap between these different clusters for some subnetworks.

A similar trend is observed in other functionally coherent assemblies of social brain regions, which are usually examined in different literature streams. For instance, the putative mirror neuron system is often thought of and studied as a cohesive neural system that includes regions like the IFG, SMG, SMA, pSTS and MTV5 (Alcalá-López et al., 2018). We found that some of these regions (e.g., the SMA) showed population co-variation with other parts of the social brain. Furthermore, these regions were not always similarly relevant in different subnetworks. For instance, subnetwork 6 featured the SMG and SMA as strong contributors together with the FP, a region which is not typically believed to be part of the canonical mirror neuron system. We thus provide evidence that widely assumed neurocognitive systems like the mirror neuron system may not prove robust to all ways to study brain-imaging data.

As another core finding that ignites future research, subnetworks 3 and 13 turned out to have predictive roles for interindividual differences in the experience of social isolation. The subjective feeling of loneliness has one of the greatest influences on some of our societies’ biggest public health concerns, in particular deep consequences for mental illness (Danilo Bzdok & Dunbar, 2020; Cacioppo & Hawkley, 2009). However, few brain-imaging studies were so far dedicated to the brain basis of perceived social isolation, which we attribute to subnetwork 13, especially the right TPJ. As one rare exception, a structural brain-imaging study found volume variability in the right TPJ to be significantly linked to rich and thin online social networks (R. Kanai et al., 2012). Based on these findings, the authors interpreted the TPJ as a region that is especially “sensitive to other people’s intentions“. Additionally, TPJ volume decline was reported in participants who self-identified as lonely (Ryota Kanai et al., 2012). These hints invite the speculation that scarcity of social interaction at home and in everyday life may reverberate in brain morphology in a way that can be quantitatively measured with common MRI scanners at the population scale.

Taken together, a few seminal studies have been dedicated to deploying clustering or latent factor methods in areas of social neuroscience. Autoencoder neural networks open the door to abstract away from clustering methods imposing strict boundaries or component discovery. In other words, at the population level, our pattern-learning technique allowed a single element of the social brain to structurally resonate with several different partner nodes. The thus extracted structural dependencies of population volume variation within our data were distinctly related to differences in social traits.

### Conclusion

We have tailored autoencoder neural networks from deep learning to perform a data-driven de-construction of an established definition of the social brain at population scale. Our fresh look into variation of structural organization suggests the existence of distinct motifs of co-dependence in these neural systems. The uncovered structural constellations of cohesive co-variation featured driving positions for the TPJ, NAC, and medial PFC. These nodes within distinct social subnetworks thus probably relate to multifaceted implementations that anchor human-defining cognitive feats, such as encoding and interrogating others’ mental states, forming social judgments, and estimating the expected value of anticipated encounters and events. Consistently, the hidden subnetwork representations, delineated by the autoencoder learning algorithms, revealed different sets of rich associations with indicators of the participants’ social capital. Many of these neurocognitive facets are traditionally studied in largely disconnected parts of the social neuroscience literature. Additionally, the revealed collection of hidden social subnetworks has potentially been overlooked by analytical approaches in widespread use. Our quantitative evidence strengthens the idea that hidden subnetworks with overlapping sources of population-level structural differentiation bring us closer to the primary biology of the social brain.

## Acknowledgments

D.B. was supported by the Healthy Brains Healthy Lives initiative (Canada First Research Excellence fund), the CIFAR Artificial Intelligence Chairs program (Canada Institute for Advanced Research), and Google (Research Award). D.B. and N.S. were also supported by NIH grant R01AG068563A.

